# LINE-1 retrotransposition impacts the genome of human pre-implantation embryos and extraembryonic tissues

**DOI:** 10.1101/522623

**Authors:** Martin Muñoz-Lopez, Raquel Vilar, Claude Philippe, Raheleh Rahbari, Sandra R. Richardson, Miguel Andres-Anton, Thomas Widmann, David Cano, Jose L. Cortes, Alejandro Rubio-Roldan, Etienne Guichard, Sara R. Heras, Francisco J. Sanchez-Luque, Maria Morell, Elisabet Aguilar, Marta Garcia-Cañadas, Laura Sanchez, Angela Macia, Pedro Vilches, Maria Concepcion Nieto-Perez, Antonio Gomez-Martin, Beatriz Gonzalez-Alzaga, Clemente Aguilar-Garduno, Adam D. Ewing, Marina Lacasana, Ignacio S. Alvarez, Richard Badge, Geoffrey J. Faulkner, Gael Cristofari, Jose L. Garcia-Perez

**Affiliations:** GENYO. Centre for Genomics and Oncological Research: Pfizer/University of Granada/Andalusian Regional Government, PTS Granada, Spain; MRC-Human Genetics Unit, Institute of Genetics and Molecular Medicine, University of Edinburgh, Western General Hospital, Edinburgh, UK; Université Côte d’Azur, CNRS, INSERM, IRCAN, Nice, France; Department of Genetics, University of Leicester, UK; The Wellcome Trust Sanger Institute, Wellcome Trust Genome Campus, Hinxton, Cambridge, UK; Mater Research Institute - University of Queensland, TRI Building, Woolloongabba, Australia; Eppendorf Ibérica, Spain; Department of Biological Geological and Environmental Sciences, University of Bologna, Italy; Agencia Sanitaria Alto Guadalquivir, Hospital Alto Guadalquivir, 23740 Andújar, Spain; Instituto de Investigación Biosanitaria Granada, Hospital Universitario de Granada-UGR, Spain; Fundacion Andaluza Investigación Biosanitaria de Andalucía Oriental Alejandro Otero (FIBAO), Granada, Spain; Department of Cellular Biology, University of Extremadura, Badajoz, Spain; Queensland Brain Institute, University of Queensland, Brisbane, Australia; Regenerative Medicine PhD Program, University of Granada, Spain; Andalusian School of Public Health (EASP), Granada, Spain; Department of Biochemistry and Molecular Biology II, Faculty of Pharmacy, University of Granada, Spain

## Abstract

Long Interspersed Element 1 (LINE-1/L1) is an abundant retrotransposon that has greatly impacted human genome evolution. LINE-1s are responsible for the generation of millions of insertions in the current human population. The characterization of sporadic cases of mosaic individuals carrying pathogenic L1-insertions, suggest that heritable insertions occurs during early embryogenesis. However, the timing and potential genomic impact of LINE-1 mobilization during early embryogenesis is unknown. Here, we demonstrate that inner cell mass of human pre-implantation embryos support the expression and retrotransposition of LINE −1s. Additionally, we show that LINE-1s are expressed in trophectoderm cells of embryos, and identify placenta-restricted endogenous LINE-1 insertions in newborns. Using human embryonic stem cells as a model of post-implantation epiblast cells, we demonstrate ongoing LINE-1 retrotransposition, which can impact expression of targeted genes. Our data demonstrate that LINE-1 retrotransposition starts very shortly after fertilization and may represent a previously underappreciated factor in human biology and disease.

## INTRODUCTION

LINE-1 retrotransposons are an abundant class of mobile genetic elements in mammalian genomes (Beck et al., 2011). In humans, LINE-1s comprise a fifth of our genomic mass and new insertions have the potential to accumulate in each passing generation (Batzer and Deininger, 2002; Beck et al., 2010; Kazazian, 1999; Lander et al., 2001). The sequencing of thousands of human genomes revealed that LINE-1s have generated millions of heritable insertions over recent evolution of the human population (Genomes Project et al., 2015; Sudmant et al., 2015). Although most LINE-1s in our genome are no longer active, an average human genome contains 80 −100 Retrotransposition Competent L1s (RC-L1s) (Beck et al., 2010; Brouha et al., 2003). As a consequence, retrotransposition is ongoing in the current human population, and new heritable LINE-1 insertions can sporadically cause genetic disease (Hancks and Kazazian, 2016; Kazazian et al., 1988). LINE-1s are also active in some somatic tissues, such as the brain (Baillie et al., 2011; Coufal et al., 2011; Evrony et al., 2012; Macia et al., 2017; Muotri et al., 2005; Upton et al., 2015) or epithelial tumors (Doucet-O’Hare et al., 2016; Ewing et al., 2015; Helman et al., 2014; Iskow et al., 2010; Lee et al., 2012; Miki et al., 1992; Rodic et al., 2015; Schauer et al., 2018; Scott et al., 2016; Shukla et al., 2013; Solyom et al., 2012; Tang et al., 2017; Tubio et al., 2014), recently reviewed in (Carreira et al., 2013; Scott and Devine, 2017)). In contrast, retrotransposition in other somatic tissues may be uncommon (Garcia-Perez et al., 2016; Macia et al., 2017). Recently, it has been demonstrated that retrotransposition in the brain can be influenced by life experience (Bedrosian et al., 2018). However, the consequences of somatic LINE-1 activity remain poorly understood, although it has the potential to contribute to tumor progression and /or tumor initiation, or even influence brain biology processes (Bedrosian et al., 2018; Doucet-O’Hare et al., 2016; Richardson et al., 2014; Rodic and Burns, 2013; Scott et al., 2016; Singer et al., 2010). At the molecular level, the detailed characterization of dozens of mutagenic LINE-1 insertions in humans illustrates how new insertions can impact the genome in a myriad of ways, from simply disrupting an exon to more complex effects on gene expression (reviewed in (Batzer and Deininger, 2002; Beck et al., 2011; Belancio et al., 2008; Garcia-Perez et al., 2016; Goodier and Kazazian, 2008; Hancks and Kazazian, 2016; Kazazian, 1999; Kazazian and Moran, 1998)). Thus, active LINE-1s are major drivers of somatic and germline genome plasticity in humans.

Despite the inherent difficulties associated with the detection of new LINE-1 insertions in genomes, estimates suggest that 1 in 100 humans and at least 1 in 8 mice might contain a *de novo* heritable LINE-1 insertion (Ewing and Kazazian, 2010; Ewing et al., 2010; Kazazian, 1999; Richardson et al., 2017; Xing et al., 2009). These vertically transmitted insertions can occur both in germ cells and during early embryogenesis, before gastrulation (i.e., in the heritable genome) (Freeman et al., 2011; Prak et al., 2003; van den Hurk et al., 2007). The use of mouse models of retrotransposition, and the sequencing of mouse pedigrees, indicate that heritable retrotransposition events occur mostly during embryogenesis (Babushok et al., 2006; Faulkner and Garcia-Perez, 2017; Kano et al., 2009; Richardson et al., 2017). In humans, rare cases of mosaic individuals carrying a pathogenic LINE-1 insertion, together with analyses of single spermatozoid genomes, strongly suggest that LINE-1 retrotransposition might also be predominant during embryogenesis (Batzer and Deininger, 2002; Brouha et al., 2002; Faulkner and Garcia-Perez, 2017; Freeman et al., 2011; Garcia-Perez et al., 2007; Kazazian, 1999; Klawitter et al., 2016; van den Hurk et al., 2007). However, the precise timing of these insertions, which cell types in the human embryo express and support LINE-1 retrotransposition, and to what extent LINE-1s drive human genomic variation remains relatively unexplored (**Figure 1a**).

**Figure 1.**
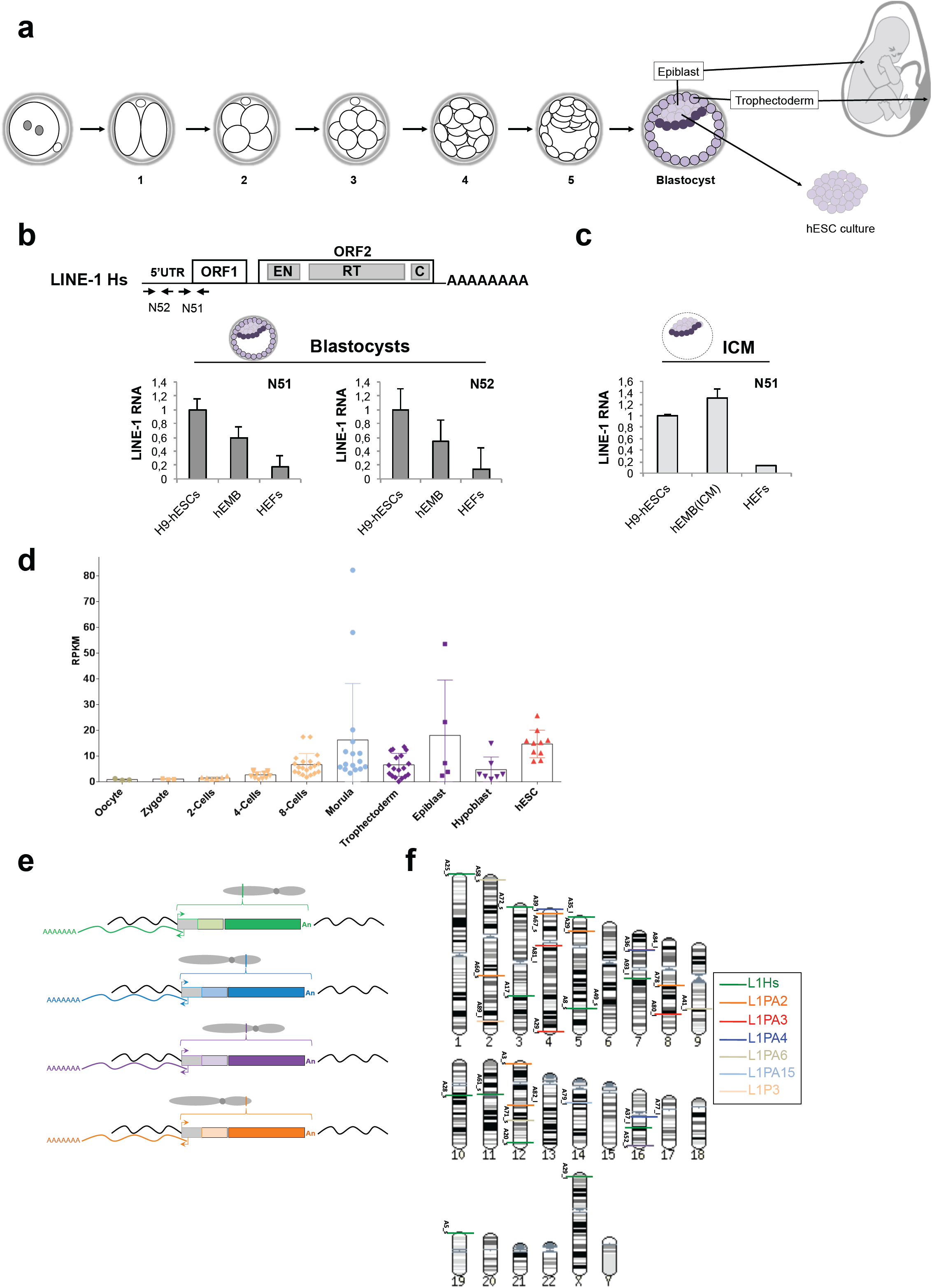
Endogenous LINE-1 mRNA expression and LINE-1-driven transcripts in human pre-implantation embryos. a, Scheme of early human embryogenesis, adapted from (Niakan et al., 2012). In the scheme, we indicate the origin of human Embryonic Stem Cells (hESCs), epiblast and trophectoderm cells. Numbers refer to embryonic days. **b**, Pre-implantation embryos (hEMBs) express L1 RNAs (total RNA extracted from a pool of 20 +5/6 day hEMBs). Shown is a schematic of a Retrotransposition-Competent LINE-1 (RC-L1); EN, Endonuclease; RT, Reverse Transcriptase; C, cysteine-rich domains. The relative position and names of primers used in RT-qPCR (black arrows) is indicated. Bar charts show the fold change in L1 RNA expression with the indicated primer pair (GAPDH normalized). Expression levels in H9-hESCs were set to 1. Error bars, standard deviation (SD) of triplicate analyses. c, H9-hESCs and the ICM of human blastocysts express similar levels of L1 RNAs. The bar chart depicts GAPDH-normalized fold change in L1 RNA expression (using Taqman) in the ICM fraction isolated from 10 human blastocysts (+6-day). Error bars, SD of triplicate analyses. d, LINE-1 RNA expression determined by RNA-seq on single cells (Yan et al., 2013). Each dot corresponds to a single-cell analyzed at the indicated stage: Oocyte (n=3), Zygote (n=3), 2-cells (n=6), 4-cells (n=10); 8-cells (n=20); Morula (n=16), Trophectoderm (n=18), Epiblast (n=5), Hypoblast (n=7), hESC (n=10). RPKM: Reads Per Kilobase per Million mapped reads. e, Identification of active antisense L1 promoters (L1-ASP, adapted from (Macia et al., 2011)). A cartoon shows 4 full-length L1s located on 4 different chromosomes. Unique transcripts generated from the L1-ASP are shown in colors: green, blue, purple and orange, and these RNAs are sequenced using a modified 3’-RACE protocol. f, L1-ASPs active in human blastocysts. The cartoon shows a human karyogram. Each cloned and sequenced L1-ASP-derived transcript identified is indicated with a horizontal colored line. Colors indicate LINE-1 subfamilies generating transcripts (L1Hs, green; L1PA2, orange; L1PA3, red; L1PA4, blue; L1PA6, light brown; L1PA15, light blue; and L1P3 light orange).

Modeling human embryogenesis with cellular models, using human embryonic stem cells (hESCs, **Figure 1a** (Thomson et al., 1998)) and human induced pluripotent stem cells (hiPSCs, (Takahashi et al., 2007)), have revealed that partially due to L1 promoter hypomethylation (Munoz-Lopez et al., 2012), a constellation of endogenous LINE-1 RNAs are overexpressed in these cells (Garcia-Perez et al., 2007; Klawitter et al., 2016; Macia et al., 2011; Marchetto et al., 2013; Wissing et al., 2011; Wissing et al., 2012). Additionally, it has been demonstrated that hESCs and hiPSCs support a moderate level of retrotransposition of engineered human LINE-1s *in vitro (*Garcia-Perez et al., 2007; Marchetto et al., 2013; Wissing et al., 2012). More recently, endogenous LINE-1s were shown to retrotranspose in cultured hiPSCs, although endogenous LINE-1 retrotransposition in hESCs has not been demonstrated (Klawitter et al., 2016). Collectively, these observations indicate that the required host factors and a permissive cellular milieu for retrotransposition are present in both hESCs and hiPSCS. However, hESCs and hiPSCs appear to mimic epiblast cells from post-implantation human embryos (i.e., after embryonic day +8) (Nakamura et al., 2016; Niakan et al., 2012). In fact, no study has analyzed LINE-1 expression or mobilization in human pre-implantation embryos (prior to embryonic day +8) (**Figure 1a**).

Here, we explored endogenous LINE-1 expression and retrotransposition in human pre-implantation embryos of the maximum quality (i.e., viable) (Cortes et al., 2007). Remarkably, we found that LINE-1 expression is detected in human pre-implantation embryos, both in Inner Cell Mass (ICM) and trophectoderm cells. Using single-cell genomics, we further demonstrate that endogenous LINE-1s are mobilized in human pre-implantation embryos. Altogether, our data demonstrate that the activity of LINE-1 elements starts shortly after fertilization, leading to genomic mosaicism in the human body, which can also be extended to extra embryonic tissues as demonstrated here (**Figure 1a**).

## RESULTS

### LINE-1 mRNA expression in human pre-implantation embryos

To examine endogenous LINE −1 expression and retrotransposition in human pre-implantation embryos (hEMBs), we screened a collection of embryos of the maximum quality available. The hEMBs used here were donated by couples who had successfully undergone IVF to produce healthy newborns (**Figure 1a**). Consistent with their presumed viability, most thawed hEMBs developed successfully to the blastocyst stage (embryonic day +6, **Figure S1**). Thus, only blastocysts with a clear and well-defined ICM were used in this study. Using RT-qPCR, we reproducibly observed abundant expression of full-length LINE-1 mRNAs in human blastocysts and hESCs (H9 line), which were used as a positive control for L1 expression (**Figure 1b and S1**) (Coufal et al., 2009; Garcia-Perez et al., 2007; Macia et al., 2017; Munoz-Lopez et al., 2012). Sequencing confirmed that ∼90% of detected L1 RNAs were derived from the L1Hs family, which encompasses all known human active L1s. Further controls confirmed that L1 ORF2 was expressed at similar levels in human blastocysts and hESCs (**Figure S2a**). Additionally LINE-1 mRNA expression was specifically observed in the ICM of blastocysts (**Figure 1c**, **S2b**). In contrast, Human Embryonic Fibroblasts (HEFs) expressed 6-10 fold less L1Hs RNA, consistent with previously described LINE-1 promoter hypermethylation in terminally differentiated cells (**Figure 1b and S2a**) (Coufal et al., 2009; Macia et al., 2017).

To extend these findings, we analysed publicly available single-cell RNA-seq data from human embryos (Yan et al., 2013). LINE-1 RNAs are present at detectable levels in oocyte, zygote, 2-cell, and 4-cell stages, with noticeable increases at the 8-cell and morula stages (**Figure 1d**). In addition, LINE-1 RNAs are readily detected in epiblast, hypoblast (i.e., primitive endoderm) and trophectoderm cells (**Figure 1d**). These data suggests that LINE-1 mRNAs are present in hEMBs prior to the initiation of Embryonic Genome Activation (EGA, (Braude et al., 1988; Dobson et al., 2004)). Similarly, we detected RNAs derived from active but non-autonomous retrotransposons at several stages of embryonic development (AluYa5 and Yb8; SVA-E, D and F) (Mills et al., 2007; Richardson et al., 2015) (**Figure S2c,d**). SVA RNAs were detected at a relatively low level up to the 4-cell stage, and became significantly elevated from the 8-cell stage forward, coinciding with EGA, and reaching maximum expression levels in morula, which considerably exceeded those observed in hESCs. Thus, these data strongly suggest that hEMBs contain high levels of L1Hs RNAs, even prior to EGA, as well as RNAs derived from potentially active classes of Alu and SVA retrotransposons.

The 5’ Untranslated Region (UTR) of LINE-1 elements contain a conserved antisense promoter (L1-ASP) (Speek, 2001) that can alter the expression of surrounding genes through alternative transcription initiation, the formation of truncated transcripts, and/or long non-coding RNA transcription (lncRNA) ((Nigumann et al., 2002), reviewed in (Garcia-Perez et al., 2016; Macia et al., 2015). Of note, sense and antisense L1 promoter activities are not necessarily coupled (Philippe et al., 2016) and recently it has been demonstrated that hundreds of L1-ASP-derived transcripts are translated in primate hiPSCs (termed LINE-1 encoded ORF0, (Denli et al., 2015)). To evaluate the potential impact of L1-ASP activity in hEMBs, we characterized transcripts initiated at the L1-ASP as previously described (Macia et al., 2011) (**Figure 1e**). We identified 58 distinct L1-ASP-derived transcripts expressed in human blastocysts. These chimeric transcripts were derived from L1 Hs elements (fixed and polymorphic), and from evolutionarily older/fixed L INE-1s, consistent with previous data from hESCs (Garcia-Perez et al., 2007; Macia et al., 2011; Philippe et al., 2016) (**Figure 1f and Table S1**). Intriguingly, we note that > 80% of identified L1-ASP transcripts expressed in hEMBs are located within annotated protein-coding genes or ESTs, consistent with a tight epigenetic control of LINE-1 expression during early development (Castro-Diaz et al., 2014; Garcia-Perez et al., 2010; Macia et al., 2011; Philippe et al., 2016; Theunissen et al., 2016). Additionally, when we compared L1-ASP RNAs expressed in hEMBs to those previously characterized in 3 hESC lines (H7, H9 and H13) (Macia et al., 2011), we identified 7 chimeric transcripts that were expressed both in hESCs and b lastocysts (**Table S1**). In particular, L1-ASP RNAs from 4p16.3, 8q21.11, 10q21.1 and 14q21.1 are expressed in blastocysts and in at least 2/3 hESCs lines, and these lncRNAs might be relevant to understanding the biology of early embryogenesis. Thus, a discrete subset of antisense LINE-1 promoters is active in blastocysts, and participates in the plasticity of the hEMB transcriptome (Macia et al., 2011; Theunissen et al., 2016).

### Detection of LINE-1 RibonucleoProtein Particles in human pre-implantation embryos

RC-L1s encode two proteins, L1-ORF1p and L1-ORF2p (**Figure 1b**), which are strictly required for retrotransposition (Moran et al., 1996). Upon translation, LINE-1 encoded proteins bind back to their encoding mRNA in a process known as *cis*-preference, forming a RiboNucleoprotein Particle (RNP) that is a presumed retrotransposition intermediate (Doucet et al., 2010; Hohjoh and Singer, 1996; Kulpa and Moran, 2005; Wei et al., 2001). In cultured cells, L1-RNPs are known to form discrete cytoplasmic foci, often enriched at the nuclear periphery (Doucet et al., 2010; Goodier et al., 2007). The formation of these cytoplasmic foci is dependent on the presence of the LINE-1 mRNA and encoded proteins, reflecting L1-RNP accumulation in cells (Doucet et al., 2010; Goodier et al., 2007).

Next, and as detecting LINE −1 RNAs does not necessarily implies that a functional LINE −1 machinery is produced, we directly examined L1-RNP accumulation in hEMBs. To do that, we used confocal microscopy and a highly specific L1-ORF1p antibody (Macia et al., 2017) (**Figure 1a**, **S3a**). As previously described, confocal microscopy analyses confirmed strong L1-ORF1p expression in hESCs, accumulating within cytoplasmic foci (**Figure 2b and S3b**) (Garcia-Perez et al., 2007; Macia et al., 2017). In hEMBs, endogenous L1-RNPs are readily detected in cytoplasmic foci, at all pre-implantation embryonic days analyzed (from day +2 to +6, **Figure 2a**). The presence of L1-RNPs indicates that a fraction of the detected LINE-1 RNAs are translated, even prior to EGA. Interestingly, at the blastocyst stage, we observed abundant L1-RNPs in ICM cells, but also in trophectoderm cells, consistent with RNA-seq data (labeled t, see **Figure 1a**, **2c**). To further validate these findings, we verified co-expression of the ICM marker POUF5F1/OCT4 and L1-ORF1p using a different L1-ORF1p antibody (Philippe et al., 2016) (**Figure S3c**). These analyses confirmed the formation of L1-RNPs in ICM and trophectoderm cells at the blastocyst stage (**Figure 2d, S3b,d**). In sum, L1-RNPs are present during early stages of human embryogenesis, even prior to EGA, consistent with parental deposition of L1-RNPs upon fertilization (Kano et al., 2009).

**Figure 2.**
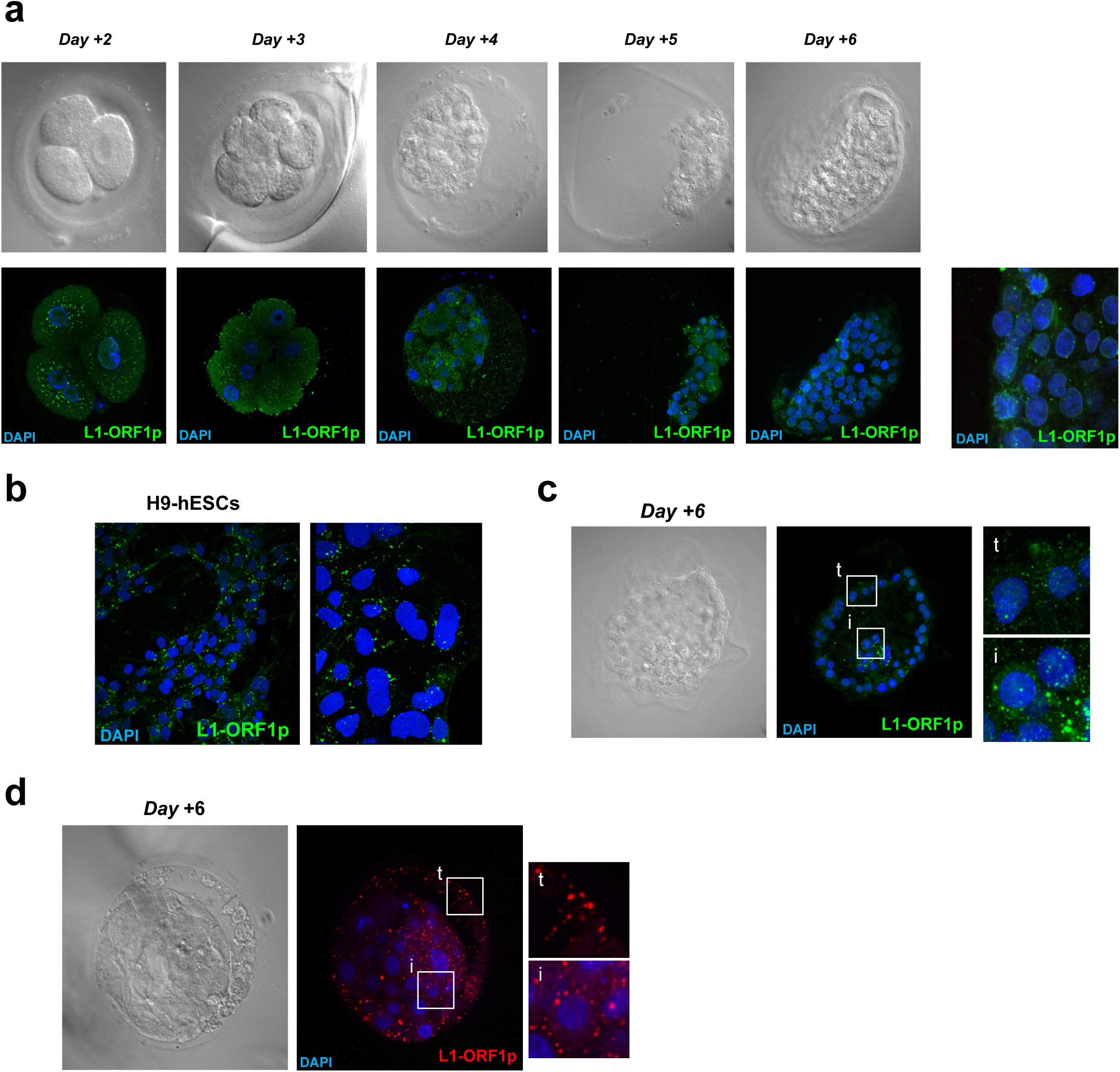
L1-RNPs are detected in human pre-implantation embryos (hEMBs). a, Shown are bright field and confocal images of hEMBs at the indicated day of development (day +2 to day +6), stained with an anti-L1-ORF1p antibody (green). Nuclear DNA was stained with DAPI (blue). Confocal pictures were captured with a 63X objective and representative merged images are shown. A magnified image of the ICM of a stained blastocyst is shown on the right. b, H9-hESCs express L1-ORF1p (as described for panel a). A magnified image is shown on the right. c and d, Trophectodermal cells express L1-ORF1p. Shown are bright field and confocal images of stained human blastocysts using two different antibodies (L1-ORF1p: green, c; red, d; nuclear DNA: blue). White boxes= magnified regions on the right (i, ICM; t, trophectoderm).

### Endogenous LINE-1 retrotransposition in the Inner Cell Mass cells of human pre-implantation embryos

The expression of L1-RNPs in hEMBs suggests that new retrotransposition events could accumulate at this stage of human development, generating genomic mosaicism in blastocysts (**Figure 3a**). To test this possibility, we applied single-cell genome analyses to hEMBs (**Figure 3a** summarizes the experimental scheme; **Text File S1** contains a full description of the procedure and of the validation of LINE-1 insertions).

**Figure 3.**
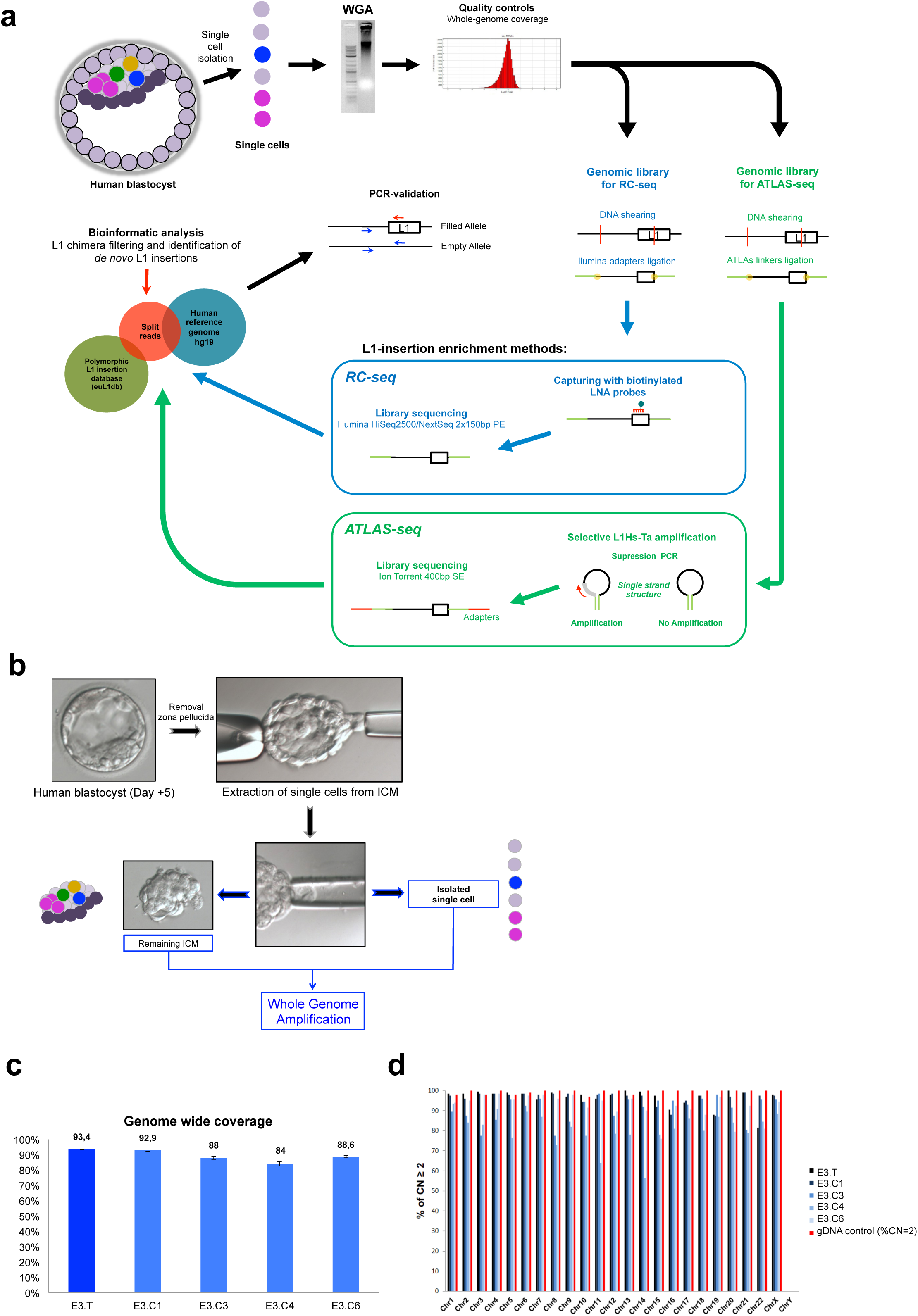
Identification of *de novo* L1 insertions in human pre-implantation embryos using single cell RC-seq and ATLAS-seq. a, Rationale of the single cell genomic approach used to identify L1 insertions in hEMBs. Whole Genome Amplification (WGA) was performed on single cells of human blastocysts. RC-seq (Sanchez-Luque et al., 2016) enrich for DNA-fragments containing L1-insertion junctions, using two biotinylated LNA (Locked Nucleic Acid) probes targeting the L1Hs-Ta consensus and exploiting diagnostic polymorphisms (Boissinot et al., 2000). Upon Illumina sequencing, bioinformatic screening detected *de novo* L1-insertions. In ATLAS-seq, suppression PCR is used to generate libraries targeting the 3’ junction L1Hs-Ta elements or the 5’-junction of full length L1Hs-Ta1d elements, using family-specific diagnostic nucleotides; libraries are sequenced by single-end 400 bp Ion Torrent (Badge et al., 2003; Philippe et al., 2016). b, Single-cell biopsies of human blastocysts. The procedure for single-cell isolation from the ICM of human blastocysts is shown, and representative pictures of the process. Isolated single cells and the rest of cells within the ICM (termed remaining embryo) were transferred to individual sterile tubes, and subjected to WGA. In the panel, ICM cells containing a specific insertion are depicted as colored circles. c, Proportion of genome amplified after WGA on samples derived from hEMB-3. An Infinium genotyping array was used (Illumina, 262,740 SNPs dispersed throughout the human genome, n=2, error bars represent SD). Locus drop out corresponds to the remaining percentage up to 100%. d, Copy number analysis of hEMB-3-derived WGA-gDNAs (data from the Infinium Assay genotyping array). The graph indicates the percentage of human genomic regions with a Copy Number (CN) of ≥2 / chromosome. Red bars, result with bulk genomic human (DNA from the blood of a healthy individual). Allele drop out corresponds to the remaining percentage up to 100%. In c and d, E3.T= WGA-gDNA from remaining embryo; E3.CX= WGA-gDNA from single cells.

Briefly, we isolated single cells from the ICM of hEMBs that reach the blastocyst stage (**Figure 1a, 3b**), amplified their genomes using Multiple Displacement Amplification (MDA, (Spits et al., 2006)), and analyzed the presence of new insertions using two different Next Generation DNA sequencing protocols: Retrotransposon Capture Sequencing (RC-Seq, (Sanchez-Luque et al., 2016)) and Amplification Typing of L1 Active Subfamilies Sequencing (ATLAS-Seq, (Philippe et al., 2016)) (**Figure 3a and Text File S1**). As validation of *de novo* LINE-1 insertions in MDA-amplified DNAs is extremely challenging (Evrony et al., 2012), we reasoned that combining two independent LINE-1 profiling methods on the same sample could help us to limit method-specific artifacts resulting from library preparation, sequencing or bioinformatic analyses (**Figure 3a and Text File S1**). Thus, we started by surgically isolating 10 single cells from the ICM of hEMBs that reached the blastocyst stage; we also collected the remaining ICM cells from each embryo and treated this heterogeneous mix of cells as an additional sample (termed *remaining embryo* or *total*, **Figure 3b, S4a**). After MDA-amplification, we only selected for further analyses th ose samples containing amplification products. Prior to library preparation, we analyzed the quality of MDA-amplified DNAs using a genomic microarray containing >250,000 SNPs (Infinium Assay from Illumina, see Methods and **Text File S1**). Using this rationale, we first analyzed cells from a hEMB that had reached the +5 days stage (termed hEMB-6), and we sequenced 2 single cells and the *remaining embryo* sample (**Figure S4a)**. After stringent bioinformatic analyses, we detected 4 putative *de novo* LINE-1 insertions in this blastocyst (**Figure 4a, left side and Tex File S1**). However, we could not completely validate any of these insertions by conventional PCR genotyping/ capillary DNA sequencing (see **Text File S1** for further details). This is not unexpected, as previous studies have demonstrated how MDA-amplified DNAs are prone to generate artifactual chimeric L1 insertions and false positives (Evrony et al., 2012; Evrony et al., 2016) (**Text file S1)**.

**Figure 4.**
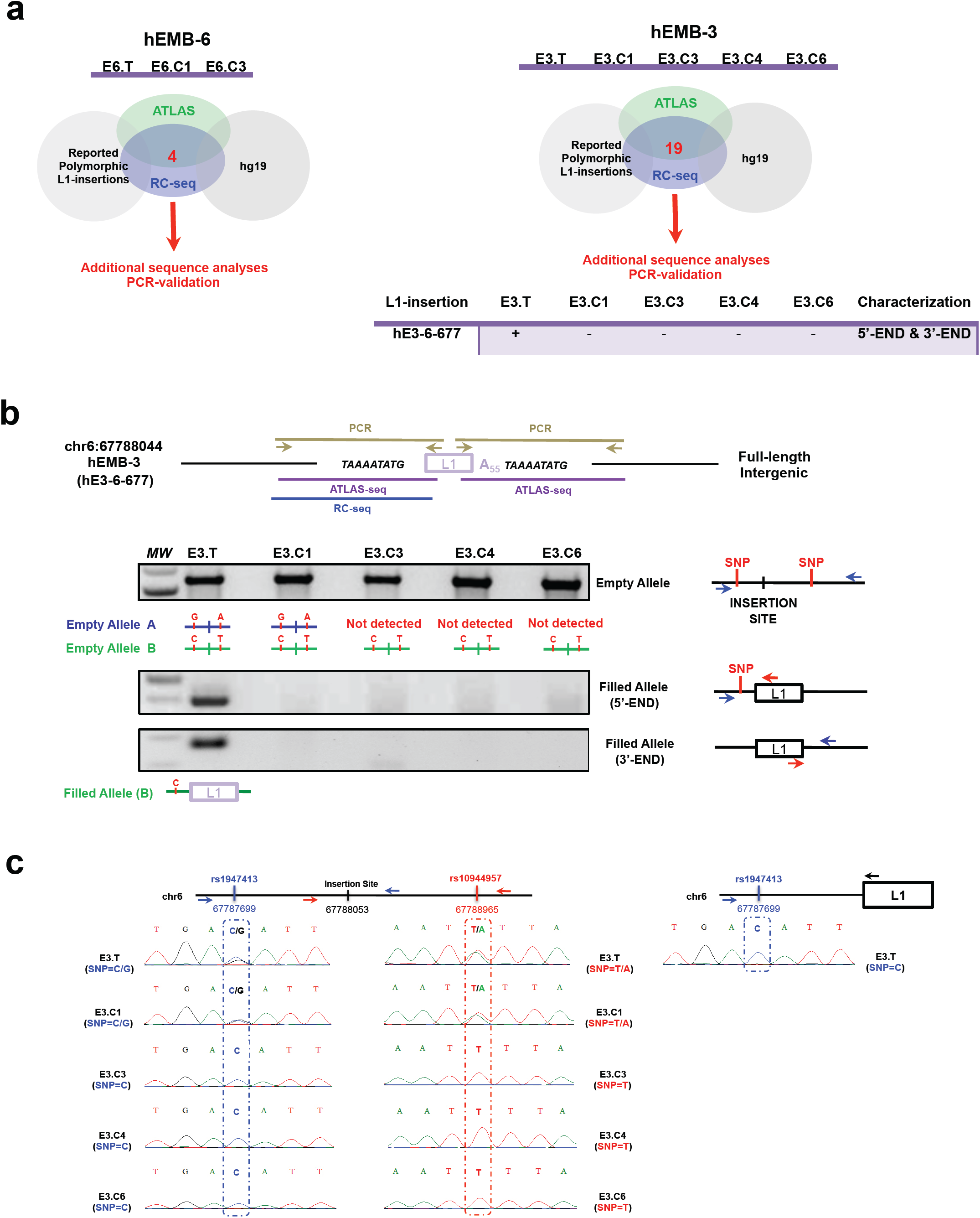
Endogenous LINE-1 retrotransposition in pre-implantation hEMBs. a, *de novo* L1-insertion candidates in hEMB-6 (left) and hEMB-3 (right). The scheme represents the screening procedure used to identify putative *de novo* L1 insertion candidates that were subsequently validated. Red colored numbers indicate the final number of insertions that were identified by RC-seq and ATLAS-seq. The table below indicates the fully validated L1-insertion by PCR/DNA-sequencing, the sample where the insertion was detected (+, present; −, absent), and the L1-ends characterized. b, validation of insertion hE3-6-677. This insertion is a full-length L1Hs-Ta insertion (chr6:67788053) identified in hEMB-3. Shown are the canonical L1-hallmarks (polyA and TSD) and the methodologies by which each end has been detected and characterized (PCR, light brown lines; RC-seq, blue lines; ATLAS-seq, purple lines). The agarose gels show representative data from genotyping PCR reactions conducted for the indicated sample (top lanes). A schematic view with the relative positions of primers (blue and red arrows) and SNPs, with respect to the insertion site, is shown. Also shown are the three allele versions of the insertion site (empty alleles A and B, and filled allele B) as detected in each sample. c, characterization of heterozygous SNPs flanking the insertion site of L1 retrotransposition event hE3-6-677. The top scheme shows the location of the insertion site and the detected heterozygous SNPs flanking the insertion site (blue and red text). Below, sequencing chromatograms from genotyping PCRs. Each detected SNP and the name of each sample are indicated (E3.T= WGA-gDNA from *remaining embryo* hEMB-3; E3.CX= WGA-gDNA from single cells isolated from hEMB-3). The right panel shows sequencing results of a PCR where we directly amplified the L1 linked to a particular SNP.

As we could not validate the putative insertions detected in hEMB-6, at least under our experimental conditions, we next analyzed an additional hEMB, but we duplicated the number of single cells sequenced. We used a hEMB that had reached the +6 stage (termed hEMB-3), and we sequenced 4 single cells and the *remaining embryo* sample (**Figure 3a, S4a)**. Analysis of concordant SNPs indicated that all selected samples were derived from the same individual (**Figure S4b**), and that high genome coverage was achieved (SNP representation in MDA-gDNAs: 84-93%) (**Figure 3c**). Furthermore, allele drop outs were evenly distributed throughout the genome, indicating that both alleles had been successfully amplified for the majority of genomic loci (**Figure 3d**). In hEMB-3, we identified 19 putative *de novo* LINE-1 insertions after RC-seq and ATLAS-seq analyses (**Figure 4a, right side**, **Text File S1** and **Table S2**). However, out of the 19 insertions, only 1 passed all the validation steps and controls performed to exclude potential artifacts or false positives (**Figure 4a, b, S5, Text File S1** and **Table S2**). This insertion (hE3-6-677) correspond to an intergenic full-length L1Hs-Ta element inserted in chromosome 6. It was validated by genotyping PCR at both termini, revealing canonical hallmarks associated with L1 retrotransposition: presence of a long polyA tail (>55 nt) and 9 bp-long Target Site Duplications (TSDs) flanking the insert ed LINE-1 element (**Figure 4b, S5**). A hemi-nested 80-cycle genotyping PCR revealed that this insertion is present in the *remaining embryo* hEMB-3 sample but not in the 4 single cells genotyped in parallel (**Figure 4b**). To exclude potential allele drop out of this *de novo* LINE-1 insertion in the 4 *single cell*s analyzed, we characterized known heterozygous SNPs located upstream (354 bp, C/G, rs1947413) and downstream (912 bp, T/A, rs10944957) of the LINE-1 insertion site by PCR and capillary sequencing (**Figure 4 b,c**). These analyses revealed that the LINE-1 insertion is genetically linked with the C SNP at rs1947413 (termed allele B in **Figure 4b, c**). Thus, any cell from hEMB-3 lacking this L1 insertion should contain the empty version of both alleles for SNP rs1947413 (alleles A and B). Remarkably, we confirmed the presence of both empty alleles in single cell E3.C1 (**Figure 4 b,c**). We could also identify the empty allele B in the rest of single cells, reinforcing that none of the single cells analysed (single cell E3.C3, .4, and .6) contained this insertion (**Figure 4 b,c**). Additionally, we confirmed the presence of allele B containing the L1 insertion, and empty alleles A and B in the *remaining-embryo* sample from hEMB-3 (**Figure 4 b,c**). Thus, hEMB-3 is a genetic mosaic with respect to this *de novo* L1 insertion (hE3-6-677, see **Text File S1** for additional validation). Altogether, these results provide proof of principle that LINE-1 retrotransposition can generate genomic mosaicism at the pre-implantation stage of human embryonic development.

### Endogenous LINE-1 expression and retrotransposition in post-implantation-like human embryonic cells

To further explore LINE-1 retrotransposition during early stages of human embryonic development, and to circumvent the inherent limitation of single-cell genomics, we next explored endogenous LINE-1 expression and retrotransposition in human embryonic stem cells (hESCs). A caveat of these analyses is the some what artifactual nature of long-term hESC cultivation, as *in vivo* these cells persist for a few days in the embryo (Niakan et al., 2012) (**Figure 1a**). However, as hESCs are more similar to post-implantation embryonic cells (Nakamura et al., 2016; Niakan et al., 2012), we reasoned that these analyses would complement the findings reported above at the pre-implantation stage (**Figure 1a**). Endogenous LINE-1 expression in hESCs has been reported before (Garcia-Perez et al., 2007; Garcia-Perez et al., 2016), but whether this is sufficient to induce endogenous LINE-1 retrotransposition is unknown (**Figure 5d, S6c**). To test this possibility, we used a well-characterized hESC line (H9 or WA09 cells, (Thomson et al., 1998)), and we analyzed a stock of H9-hESCs that has been cultivated in our lab for the past 10 years (with routine quality controls and monitoring of passage number). We confirmed that our stock of H9-hESCs overexpress LINE-1 mRNAs (**Figure 5a, S6a**) and L1-ORF1p (**Figure 5b, S6d**), and that L1-RNPs are readily detected in POUF5F1/OCT4 expressing cells, as cytoplasmic foci (**Figure 5c, S3b**). Next, and to explore LINE-1 retrotransposition, we expanded a culture of H9-hESCs (starting at passage 58, with a stable karyotype, **Figure S6f**) for 10 consecutive passages, harvesting genomic DNAs during the process. Next, we applied RC-seq to H9-hESCs at passages p58 and p69 (Sanchez-Luque et al., 2016; Upton et al., 2015), using high molecular weight (HMW) DNAs (**Figure 5d**). Upon bioinformatic analyses to remove annotated and polymorphic L1Hs insertions (see **Text File S2** for additional details on validation of insertions in hESCs), we validated and characterized a *de novo* LINE-1 retrotransposition event (H9-16-723) (**Figure 5d-g**). Insertion H9-16-723 is a nearly full-length L1Hs-Ta element inserted in intron 11 of LINC01572, a long non-coding RNA gene located on chromosome 16, in the sense orientation (with respect to LINC01572, **Figure 5g**). The insertion was fully characterized at both ends, revealing the presence of canonical hallmarks of retrotransposition (**Figure 5g, S7a**). Validation demonstrated that, while the empty site is detected in all passages tested (p58, p60, p67 and p69), L1-insertion H9-16-723 was detected only at p67 and p69 (**Figure 5f**), suggesting that retrotransposition occurred during the culturing of H9-hESCs.

**Figure 5.**
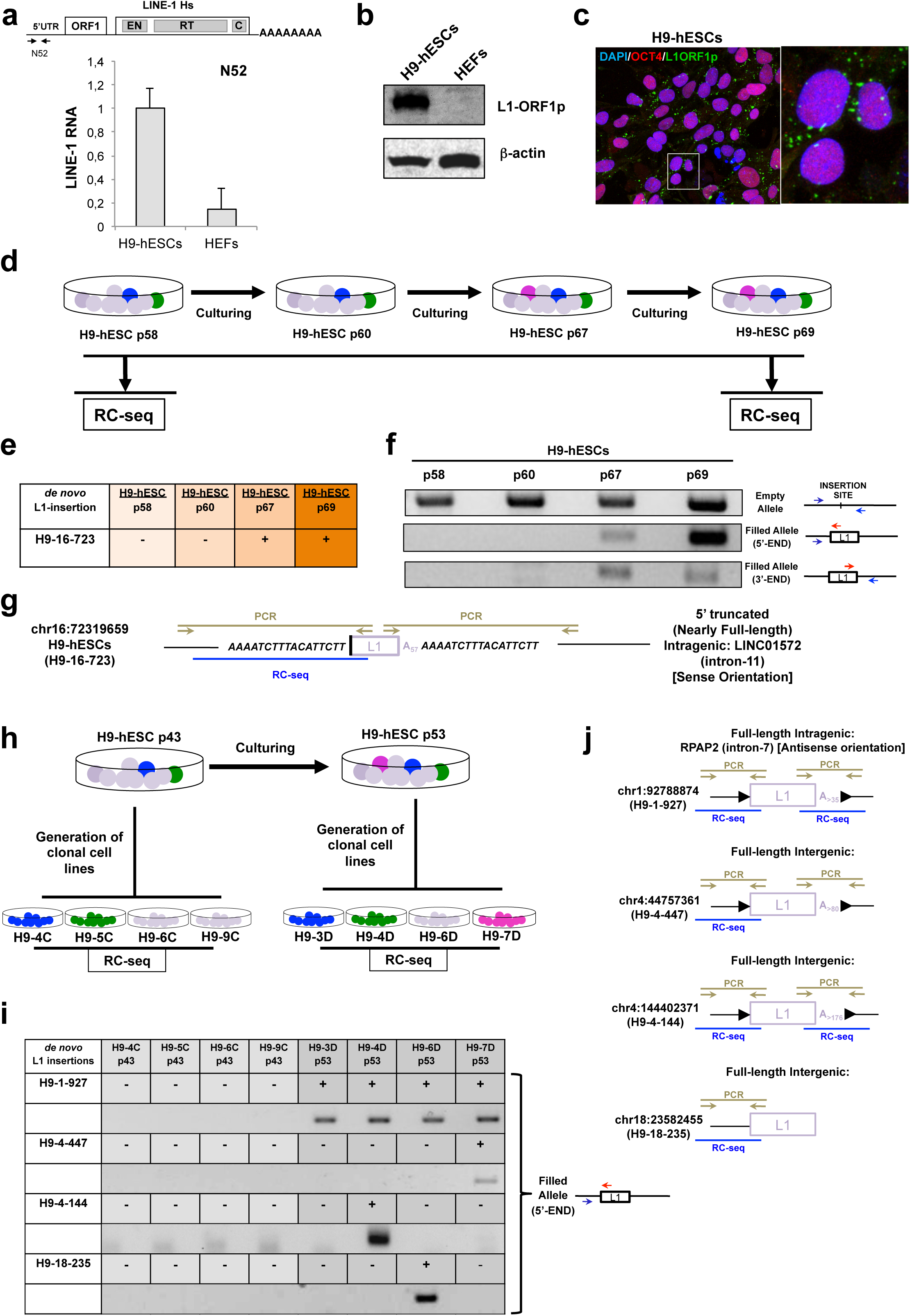
Endogenous LINE-1 expression and retrotransposition in human embryonic stem cells. a, hESCs express LINE-1 mRNAs. Schematic of an active L1, where the relative position and name of the primer pair use d in the RT-qPCR is shown (N52, small black arrows). H9-hESCs expression was set to 1. Error bars, SD of triplicate analyses. b, hESCs express L1-ORF1p. Western-blot assay on whole cell lysates from H9-hESCs and HEFs (see Figure S3a for antibody characterization). β-actin was used as a loading control. c, hESCs express L1-RiboNucleoprotein Particles (L1-RNPs). Shown is a merged image of H9-hESCs stained for L1-ORF1p (green) and POUF5F1/OCT4 (red) expression; nuclear DNA was stained with DAPI (blue). Confocal pictures were captured with a 63X objective; white square = magnified section of the image shown on the right. d, Rationale of RC-seq experiments conducted on hESCs grown as pools. New L1 insertions generated upon culturing hESCs are depicted as colored circles. H9-hESCs were cultured for 69 passages, and genomic DNAs prepared at passages 58, 60, 67 and 69. The profile of L1 insertions at passages 58 and 69 was compared. e, Validation of *de novo* L1-insertions characterized on pools of H9-hESCs. The + sign indicates if the identified *de novo* L1 insertion (insertion H9-16-723) was detected by PCR at the indicated passage. f, Agarose gel show ing representative data from genotyping PCR reactions for the empty site and the 5’ and 3’ ends of insertion H9-16-723. A schematic with the relative positions of primers (blue and red arrows) is shown. g, Schematic of insertion H9-16-723. Nearly full-length (truncated at nucleotide 21 with respect to L1.3 (Sassaman et al., 1997)) that inserted into intron 11 of the long non-coding RNA LINC01572, sense orientation. Shown are the canonical LINE-1 retrotransposition hallmarks (a long polyA tail (>57 nt) and 17 bp-long TSDs) and the methodologies by which each end was detected and characterized (PCR, light brown lines; RC-seq, blue lines). h, Rationale of RC-seq on clonal sublines established from hESCs at passages 43 (left) and 53 (right) (see Methods and Figure S6h). New L1 insertions generated upon culturing hESCs (colored circles) can be segregated in the clonal sublines. i, RC-seq results on clonal sublines. The +/- signs indicate if the identified *de novo* L1 insertions are present in clonal cell lines. Embedded are shown agarose gels with representative data from genotyping PCR (a schematic with the positions of primers (blue and red arrows) is also shown). j, Schematic of insertions validated in clonal sublines. Shown are the canonical L1 retrotransposition hallmarks (long polyA tails (from 35 to >176 nt) and flanked by TSDs (14 to 18 bp-long)) and the methodologies by which each insertion has been detected and characterized (PCR, light brown lines; RC-seq, blue lines). Also indicated is the location of each insertion.

The above experiments demonstrate that endogenous LINE-1s can retrotranspose in the genome of post-implantation-like embryonic cells, although the identification of new insertions is challenged by the heterogeneity of cultures. To overcome this limitation, we generated and expanded clonal cell lines from cultures of H9-hESCs (**Figure S6c**). We started with a heterogeneous culture of H9 - hESCs with a normal and stable karyotype (at p43 (**Figure S6g**)), and we isolated 8 clonal cell lines (4 at passage p43 (**Figure 5h, left side, S6h**) and 4 at passage p53 (**Figure 5h, right side, S6h**)). Then, we conducted RC-seq on HMW-genomic DNAs isolated from the 8 clonal lines (**Figure 5h**). After bioinformatic screening (**Text File S2**), we validated and fully characterized 4 *de novo* LINE-1 insertions differentially present among the analyzed clones (**Figure 5i-j, S7d**). These four insertions (H9-1-927, H9-4-447, H9-4-144 and H9-18-235) are all full-length L1Hs-Ta elements and exhibit canonical retrotransposition hallmarks (**Figure 5j, S7d**). Three insertions occurred into intergenic regions, while insertion H9-1-927 occurred intron 7 of the *RPAP2* gene, in antisense orientation (**Figure 5j, S7d**). While the empty site of the four *de novo* insertions could be amplified in the 8 clonal cell lines (**Figure S7b**), insertion H9-1-927 was detected in the 4 subclonal cell lines isolated at H9-hESC-p53 (**Figure 5i**), strongly suggesting that it occurred between passages p43 and p53. In contrast, insertions H9-4-447, H9-4-144 and H9-18-235 were only present in the sublines H9-7D-p53, H9-4D-p53 and H9-6D-p53, respectively (**Figure 5i**). These data may suggest that these three retrotransposition events accumulated later, and were present in a mosaic manner in the H9-hESC culture used to derive the sublines (H9-hESC-p53), although we cannot rule out that these insertions occurred during the cloning or culturing of the clonal cell lines.

Next, we selected clonal subline H9-4C for further analyses; this clonal subline was established from H9-hESC-p43 and no *de novo* LINE-1 insertion was detected in this clonal cell line when compared to the other 7 H9-hESC-derived sublines analyzed (**Figure 5**). Quality controls of this subline at passage 0 (H9-4Cp0) revealed presence of a normal and stable karyotype (**Figure 6a**), strong alkaline phosphatase activity (**Figure S7f**), expression of pluripotent markers (TRA1-60 and POUF5F1/OCT4, **Figure 6b**) and the capability to differentiate into the main three germ layers in a teratoma assay (**Figure 6c**) or using an Embryoid Body (EB)-based differentiation assay (**Figure S7g**). In sum, this clonal subline behaves the same as the parental cell line H9-hESC. Similar data were obtained with other characterized H9-derived sublines. Having phenotypically and genetically characterized the H9-4C subline, we cultured it for 10 additional passages, and conducted RC-seq (on HMW-DNAs, **Figure 6d**). After bioinformatic screenings, we validated and characterized three *de novo* LINE-1 insertions present at passage p10 but absent at passage p0 (**Figure 6d-f, S7c and S7e**), suggesting they occurred while culturing this subline. The three insertions (H9-4C-12-192, H9-4C-3-115 and H9-4C-21-369) are full-length L1Hs-Ta elements, show canonical retrotransposition hallmarks, and are inserted into known genes (**Figure 6f** and **S7e**, 12-192, in intron 26 of gene *CACNA2D4*, in antisense orientation; 3-115, in intron 3 of gene *LSAMP*, in sense orientation; and 21-369, in intron 5 of gene *RUNX1*, in antisense orientation). Similarly, culturing a second subline for 14 passages (H9-4D, derived from H9-hESC-p53), led to the identification of another *bona fide* LINE-1 insertion, as revealed after RC-seq analyses (**Figure S8a**). This insertion is present at p14 and absent at p0 (**Figure S8b-d**), yet again is full-length and inserted in intron 1 of the *KRTAP5-AS1* gene, in antisense orientation (**Figure S8c-d**).

**Figure 6.**
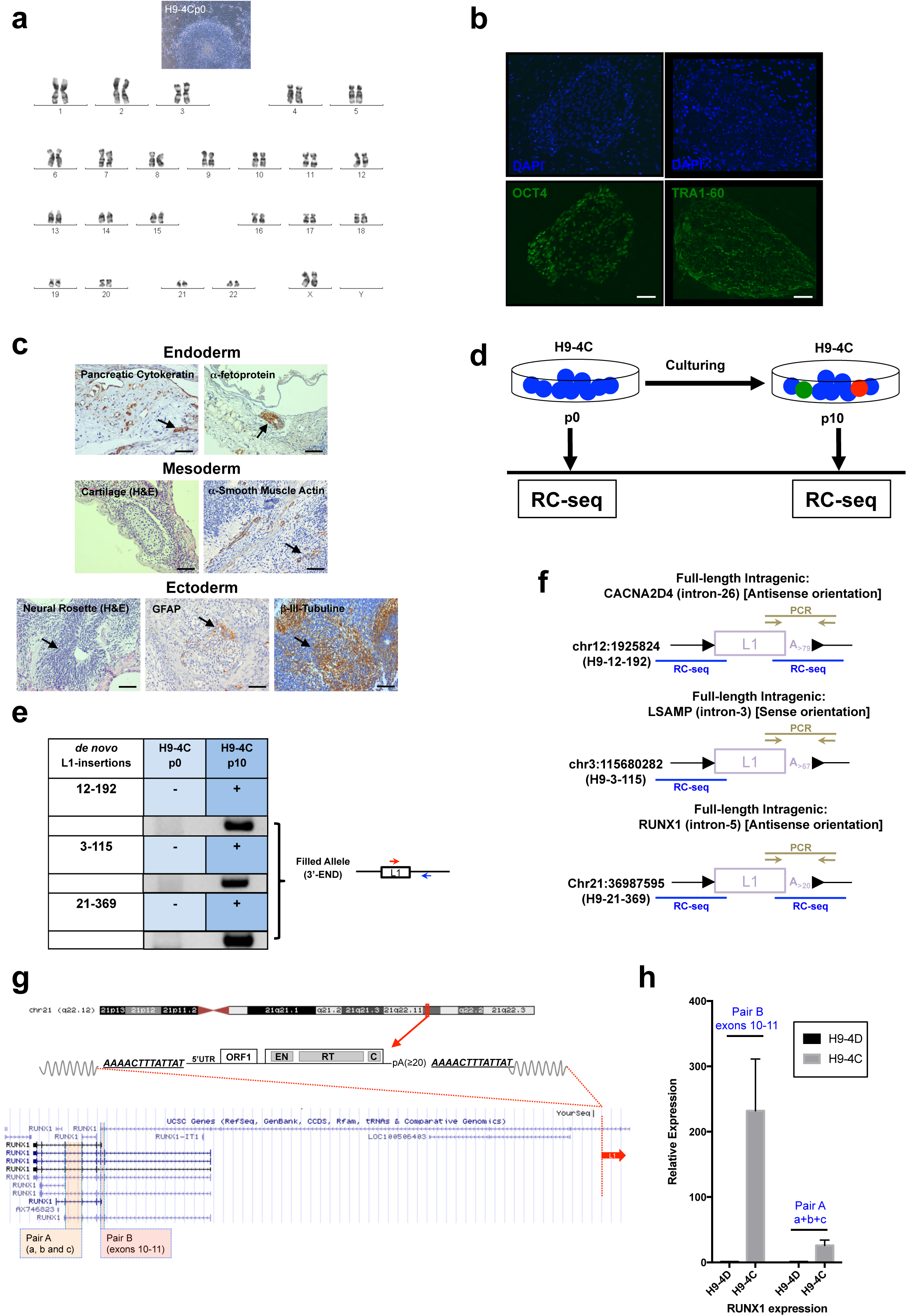
Genic LINE-1 retrotransposition events can impact gene expression. a, Representative light microscopy picture of H9-4C cells and karyogram at passage 0. A representative G-banding metaphase staining is shown (from thirty captured). b, H9-4C cells express pluripotent markers. H9-4Cp0 cells were stained with POUF5F1/OCT4 (green, left side) or TRA1-60 antibodies (green, right side); nuclear DNA was stained with DAPI (blue). White scale bars, 100∞m. c, Teratoma assay using H9-4Cp0 cells. Shown are results from Hematoxylin & Eosin (H&E) staining, and of staining with Pancreatic Citokeratine and α-fetoprotein (Endoderm), α-smooth-muscle actin (Mesoderm), and GFAP (Glial Fibrillary Acidic Protein) and β-III-tubulin (Ectoderm). Black arrows= positive foci. Scale bars, 500∞m. d, Rationale of the RC-seq analyses on the clonal subline H9-4C (expanded for 10 passages). New L1 insertions are shown as colored circles. e, RC-seq results on subline H9-4C. The +/- signs indicate the presence of *de novo* L1 insertions. Embedded in the table are shown agarose gels with representative data from genotyping PCR reactions (a schematic with the positions of primers (blue and red arrows) is also shown). f, Schematic of insertions validated in expanded clonal sublines. Shown are the canonical L1 retrotransposition hallmarks (poly A tails (from 20 to >79 nt) and TSDs (from 13 to 17 bp)) and the methodologies by which each insertion has been detected and characterized (PCR, light brown lines; RC-seq, blue lines). Also indicated is the location of each insertion. g, Genomic environment of insertion H9-4C-21-369. Cartoon of chromosome 21, where the insertion site is indicated with a red box. A cartoon of this full-length insertion is shown below (TSDs, uppercase italic lettering). Also shown are the different isoforms of RUNX1 from UCSC Genome Browser (hg19) and the relative position of the two primers pairs (Pair A and B) used to analyze RUNX1 expression by RT-qPCR. Pair A (orange shading) detects isoforms a, b, and c among others. Pair B (red shading) detects isoforms spanning the site of this LINE-1 insertion, which is intron 5. h, RT-qPCR RUNX1 expression analyses. The graph indicates the relative expression (GAPDH normalized) of RUNX1 in clonal sublines that contain (H9-4C, grey bars) or lack (H9-4D, black bars) this insertion. Indicated in the graph is the primer pair used (blue text) and the SD of triplicate analyses.

Importantly, additional controls revealed that the nine LINE-1 insertions characterized in this study on H9-hESCs are absent from the genome of other stocks of these cells grown elsewhere (i.e., as demonstrated after analyzing the genome of a H9-hESCs stock sequenced within ENCODE (**Text File S2**)). In summary, the analyses conducted in H9-hESCs, a model of post-implantation embryonic cells, revealed that endogenous LINE-1s retrotranspose efficiently in these cells. LINE-1 insertions were validated in pools of heterogeneous cultures, clonal sublines, and cultured clonal sublines, leading to the accumulation of LINE-1 driven genomic mosaicism in these cells.

### Genic endogenous LINE-1 retrotransposition events in post-implantation embryonic-like cells can impact gene expression

Notably, all LINE-1 insertions we validated in pluripotent cells were full-length (n=10, 1 in hEMBs, and 9 in hESCs). The accumulation of only f ull-length LINE −1 insertions is unexpected (see Discussion and **Figure S19**). To further exclude that RC-seq preferentially identifies full-length LINE-1 insertions, we next analyzed in parallel two lung cancer HMW-genomic DNAs, as retrotransposition events have been characterized in this type of human tumor (Iskow et al., 2010). To note, using the same RC-seq protocol, all PCR/DNA-sequenced validated *de novo* LINE-1 insertions in lung tumors were 5’-truncated (n=10, **Figure S19**). Thus, in principle our study has no bias toward the identification of full-length LINE-1 insertions.

On the other hand, two thirds of the new retrotransposition events validated in H9-hESCs occurred into known human genes (6 out of 9). Thus, we next analyzed the impact of genic LINE-1 insertions on the expression of their target gene: *RUNX1 (*insertion present in subline H9-4C, at p10); *RPAP2 (*insertion present in sublines H9-3D, H9-3D, H9-6D and H9-7D at p0); *LSAMP (*insertion present in subline H9-4C, at p10); and *LINC01572 (*insertion present in a pool of H9-hESCs at p69). *RUNX1* (Runt Related Transcription Factor 1) is a transcription factor involved in human development and key for hematopoiesis; in addition, it can act as an oncogene (reviewed in (Sanda, 2017)), and chromosomal translocations involving *RUNX1* are common in several types of leukemia (reviewed in (de Bruijn and Dzierzak, 2017)). *RPAP2 (*RNA Polymerase II Associated Protein 2) is involved in gene transcription and specifically regulates small nuclear RNA (snRNA) transcription (Jeronimo et al., 2016). *LSAMP (*Limbic System-Associated Membrane Protein) has been involved in the development of the nervous system, mediating neuronal growth and exon targeting (Eagleson et al., 2003). As a lncRNA, *LINC01572 (*Long Intergenic Non-Protein Coding RNA 1572) could be involved in a myriad of biological processes, from gene expression to stem cell maintenance (Moran et al., 2012). The comparison of H9-hESCs clonal sublines that differentially contain or lack these insertions in genes *RPAP2, LSAMP* and *LINC01572*, revealed only minor and non-significant changes in gene expression induced by the new LINE-1 insertion (<0.5-fold differences), at least under experimental conditions (**Figure S9a-c**).

In contrast, insertion 21-369 occurred in the *RUNX1* gene (intron 5, antisense insertion) and coincided with significant changes in gene expression (**Figure 6 g,h**). Indeed, we observed a large and significant increase in *RUNX1* RNA expression in H9-4C p10 cells carrying the insertion when compared to cells lacking the insertion (H9-4D line, 231 ± 79 fold; **Figure 6g**, pair B, **6h**). There are more than 15 distinct annotated *RUNX1* transcripts in Ensembl, and three of them are well-known characterized isoforms which are termed a, b and c (Sanda, 2017) (**Figure 6g**). Thus, we next analyzed whether the intronic *RUNX1* LINE-1 insertion induced changes in gene expression in the a, b and c isoforms of *RUNX1 (*using a second set of primers, Pair A, **Figure 6g**). Consistently, we observed that hESCs carrying the insertion (H9-4C) overexpress *RUNX1* transcripts (26 ± 8 fold; **Figure 6h**), including the a, b and c isoforms of *RUNX1.* A similar increase in RUNX1 expression was detected when gene expression levels were compared to other sublines lacking the insertion (i.e., H9-7D, data not shown). In sum, these data show that insertion H9-21-369, which is specifically present in the H9-4C subline, results in the overexpression of *RUNX1*, which could impact the fate of these hESCs (see Discussion).

### Endogenous LINE-1 retrotransposition in extraembryonic tissues

So far, our results strongly suggest that LINE-1 retrotransposition can take place in pre-implantation embryos, and likely in post-implantation pluripotent cells, ultimately leading to L1-driven genetic mosaicism in the adult human body. On pre-implantation embryos, we also detected abundant L1-derived RNAs and RNPs in trophectoderm cells of blastocysts (**Figure 1d, 2**), which contribute to extraembryonic tissues and placenta formation (**Figure 1a**). These expression data is consistent with previous reports documenting LINE-1 RNA expression in placental tissues (Ergun et al., 2004; Reichmann et al., 2013). To test if LINE-1 expression in this cell type could result in the accumulation of new extraembryonic-restricted LINE −1 insertions, we collected extraembryonic tissues samples from two newborns and analyzed the profile of LINE-1 insertions using RC-seq (**Figure 7a**, using HMW-genomic DNAs isolated from newborn-1 and −2, (Sanchez-Luque et al., 2016)). Specifically, we collected the following samples from newborns: amnion (derived from epiblast), chorion leave and chorion frondosum (from polar trophectoderm) and newborn blood, as well as maternal blood genomic DNA as a control (**Figure 7a**). Notably, we validated and fully characterized one *de novo* LINE-1 insertion in these samples (n1-5-174, **Figure 7b, S10a and Table S6**). Insertion n1-5-174 is an intergenic full-length L1Hs-Ta insertion on chromosome 5, with canonical hallmarks of retrotransposition (**Figure 7b, S10a**). This insertion was present only in the chorion frondosum (that is the embryo-derived placenta), and was found absent in all other samples from newborn-1, despite the use of 80-cycles nested-PCR (**Figure 7b, S10a and Table S6**). We speculate that this insertion could have accumulated in the polar trophectoderm during early embryogenesis, consistent with the pattern of L1 expression on blastocysts.

**Figure 7.**
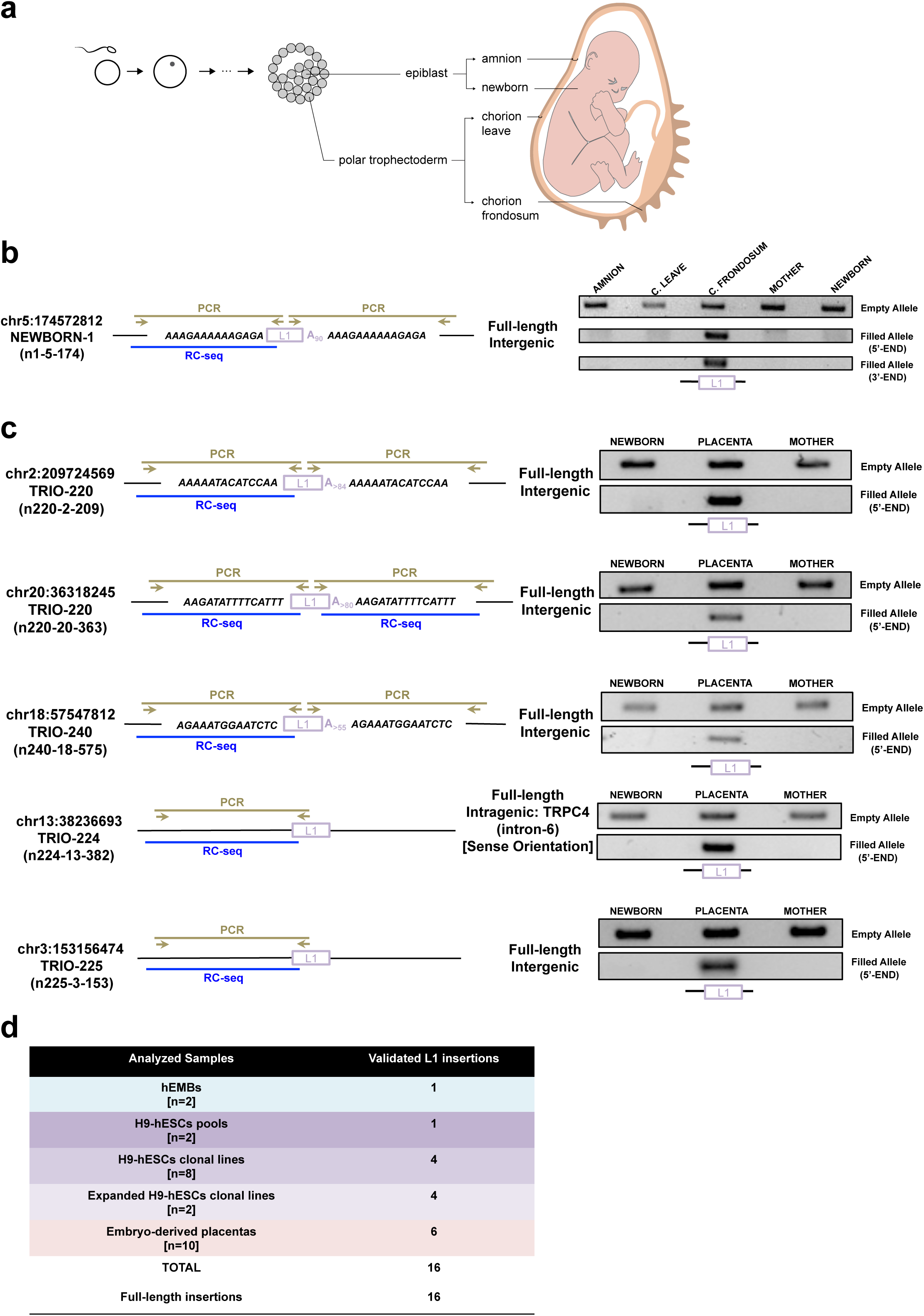
LINE-1 retrotransposition in extraembryonic tissues. a, The schematic representation depicts a newborn with its extraembryonic tissues as well as their embryonic origin from the blastocyst stage. Chorion leave and chorion frondosum (embryo-derived placental tissue) derive from polar trophectoderm cells; amnion and newborn tissues are derived from the epiblast. HMW-DNAs from these tissues were analysed by RC-seq; >90% of refere nce L1Hs elements were recovered in these RC-seq libraries (hg19). b, Validation of insertion n1-5-174. This LINE-1 insertion is a full-length L1Hs-Ta element flanked by 14-bp long TSDs and terminating in a polyA tail >90 residues in chromosome 5 (position 174572812). The right side shows representative data from genotyping PCR reactions for the empty allele (top gel), the 5’-filled end (middle gel) and the 3’-filled end (bottom gel). Sample name is indicated above each lane: amnion tissue (AMNION), chorion leave (C. LEAVE), chorion frondosum (C. FRONDOSUM), mother’s blood (MOTHER) and newborn’s blood (NEWBORN). c, Validation of insertions n220-2-209, n220-20-363, n240-18-575, n224-13-382 and n225-3-153. As in panel b, the right side shows representative data from genotyping PCR reactions, for the empty allele (top gel) and the 5’-filled end (bottom gel). The sample is indicated above each lane: embryo-derived placenta (PLACENTA), mother’s blood (MOTHER) and newborn’s blood (NEWBORN). In b-d, the hallmarks associated with L1 retrotransposition and the detection methods are shown (PCR, light brown lines; RC-seq, blue lines). Note that canonical hallmarks of retrotransposition could only be annotated in fully validated insertions. d, Summary of the 16 *de novo* L1 insertions characterized in this study.

To further explore the frequency of LINE-1 retrotransposition in placental tissues, we next analyzed eight additional conceptions by RC-seq, comparing embryo-derived placenta, newborn cord blood, and maternal blood (HMW-genomic DNAs were extracted from 4 pooled placental sections, see Methods). Upon bioinformatic screening, we selected a group of putative *de novo* LINE-1 insertions present only in the placenta of the 8 newborns analyzed (**Table S6**), and 5 were validated (n220-2-209, n220-20-363, n224-13-382, n225-3-153 and n240-18-575, **Figure 7c, S10b and c**). Remarkably, the 5 insertions were full-length L1Hs-Ta elements, and two were present in the same placenta (newborn n220) (**Figure 7c and S10b, c**). Additionally, one of these insertions, n224-13-382, was inserted into intron 6 of gene TRPC4, in antisense orientation. Overall, we have confirmed a *total* of 6 *de novo* LINE-1 insertions in the embryo-derived placentas of 5 out of 10 newborns (50%), demonstrating that L1-driven genetic heterogeneity can also impact extraembryonic lineages, resulting in the accumulation of non-heritable L1 insertions.

## DISCUSSION

The genomic revolution provides a unique opportunity to understand the genomic diversity of the human body, especially using single-cell genomics. In this study, we exploited high-throughput sequencing data, as well as cell imaging, to analyze endogenous LINE-1 expression and retrotransposition at several time points during human embryonic development (**Figure 1a, 7d**). At the RNA level, LINE-1 transcripts accumulate during early embryonic development, from embryonic day 0 to +6 (**Figure 1**). Additionally, we have characterized a specific class of lncRNAs present in pre-implantation hEMBs and that are transcribed from the antisense promoter of full-length LINE-1s (L1-ASP, **Figure 1f**). In particular, we found that a small group of L1-ASP derived lncRNAs are expressed in hEMBs and most hESCs examined (loci 4p16.3, 8q21.11, 10q21.1 and 14q21.1, **Table S1**), and could be candidate biomarkers of pluripotent cells, as proposed recently for other TE-transcribed loci (Theunissen et al., 2016). Thus, LINE-1s participate in the diversification of the transcriptome of human pre-implantation embryos. Next, by analyzing single cells, we demonstrate that both the expression and ongoing mobilization of endogenous LINE-1s can occur in the ICM fraction of pre-implantation human embryos (**Figures 2-4**). From a technical angle, we found that the combination of two distinct LINE-1 profiling methods significantly increases the successful identification and validation of *de novo* LINE-1 insertions when examining MDA-amplified DNAs (**Text File S1**). From a biological angle, we validated one full-length LINE-1 retrotransposition in ICM cells after sequencing just 6 single cells and 2 remaining ICMs from two different hEMBs. This hEMB insertion occurred very early during embryogenesis, and could have the potential to be transmitted to the next generation, as germ cells appear at a much later stage. Although we cannot pinpoint the precise timing of retrotransposition, the absence of this insertion in all single cells analyzed from hEMB-3 suggests that the retrotransposition event occurred post - fertilization, and after a few rounds of cell division. Additionally, non-autonomous but active retrotransposons belonging to the SVA or Alu families are highly expressed in hEMBs, and may utilize the LINE-1-encoded machinery to mobilize at this stage of development (**Figure S2**). Altogether, these data demonstrate that *de novo* heritable LINE-1 insertions can accumulate during early human embryogenesis, prior to embryo implantation. Consistent with this, in a recent study in mouse, where we followed LINE-1 retrotransposition in mouse pedigrees, we demonstrated that the activity of retroelements in the heritable genome gives rise to a patchwork of distinct genetic territories in the mammalian body (Richardson et al., 2017). Thus, despite differences in the overall structure and sequences of human and mouse LINE-1s, as well as marked differences in early embryonic development programs, these data suggest that retrotransposition during early embryogenesis is conserved in mammals. With respect to the frequency of LINE-1 retrotransposition in pre-implantation hEMBs, it is noteworthy that due to the sensitivity of MDA-single-cell sequencing and the conservative approach chosen to validate *de novo* L1 insertions in this study, it is likely that what we are reporting here represent just the “tip of the iceberg”. Moreover, as we only compared a small number of single cells from pre-implantation human embryos, very early retrotransposition events will be present in most/all cells of a given blastocyst, and thus will not be identified as putative *de novo* L1 insertions following the experimental approach of our study. We speculate that the rate of LINE-1 retrotransposition in ICM cells at the pre-implantation stage of development could be significantly higher than reported here. The quality of the embryos used in this study is excellent, as we used embryos that were sibling of babies that were born after a successful round of IVF (Cortes et al., 2007). Whether IVF increases the rate of endogenous L1 retrotransposition in pre-implantation embryos requires further elucidation.

The post-implantation stage in humans starts around day +8 (Niakan et al., 2012), and although hESCs are derived from the ICM of pre-implantation embryos, recent studies have demonstrated that cultured hESCs resemble pluripotent stem cells present at the early, post-implantation stage of human development (Nakamura et al., 2016). Intriguingly, several lines of evidence suggest that LINE-1 retrotransposition can also occur later during embryonic development. Indeed, studies in mouse have demonstrated the accumulation of LINE-1 insertions during later development, resulting in L1 insertions that are present only in selected somatic tissues of the body and cannot be transmitted to the next generation (Kano et al., 2009). Thus, in this study we used hESCs as a model of post-implantation pluripotent cells of the human embryo, building on previous reports describing endogenous LINE-1 expression in hESCs and other pluripotent stem cells (Garcia-Perez et al., 2007; Garcia-Perez et al., 2010; Klawitter et al., 2016; Marchetto et al., 2013; Wissing et al., 2011; Wissing et al., 2012). To this end, we followed the accumulation of new LINE-1 insertions during the culturing of hESCs (i.e., examining pools at different passages), but also when clonal hESC sublines are generated and expanded (**Figure 5 and 6**). We uncovered 9 *de novo* LINE-1 insertions in hESCs, and remarkably 6 out of 9 (67%) inserted into the introns of known protein-coding genes or a lncRNA. Most of these genic LINE-1 insertions had minor effects in gene expression (for *RPAP2, LSAMP* and *LINC01572*), at least at the hESC stage. However, one genic insertion into the *RUNX1* gene led to a large and significant overexpression of *RUNX1* RNAs in cells carrying this new LINE-1 insertion. A possible mechanism could involve the LINE-1 antisense promoter and alternative transcription initiation, as this promoter is very active in pluripotent cells (Denli et al., 2015; Macia et al., 2011; Nigumann et al., 2002) (**Figure 1f**). However, there are several alternative mechanisms that could be involved in *RUNX1* deregulation, as alterations of regulatory sequences and splicing patterns, or epigenetic perturbations induced by the insertion. As *RUNX1* is a key transcription factor for hematopoiesis, often deregulated in leukemia, it is very likely that the fate of hESCs carrying this insertion would be affected. Altogether, here we demonstrate that LINE-1s impact the genome of post-implantation embryonic-like cells, and that the impact in gene expression is variable and likely influenced by the site, type and length of the insertion, the orientation of the insertion, and the cell type analyzed, among other factors. Whether the accumulation of new LINE −1 insertions in hESCs/hiPSCs can limit the biomedical potential of these cells should be explored in future studies (Klawitter et al., 2016).

Intriguingly, we also found that L1-RNPs are detected in trophectodermal cells of pre-implantation hEMBs. Thus, we exploited high-throughput sequencing to analyze whether LINE-1s retrotranspose in extraembryonic tissues, such as the embryo-derived placenta. After examining 10 newborns, we demonstrated that retrotransposition in placental tissues is common, and 50% of the examined placentas had at least one *de novo* L1 retrotransposition event. It is likely that this percentage might be higher (perhaps 100%), as we only analyzed a small fraction of such large tissue, and our study is biased to the detection of very early retrotransposition events. Because of their early embryonic origin, for any insertion characterized in placenta, we cannot exclude that these insertions might also be present in other tissues from newborns, although this is very unlikely. Indeed, in stark contrast to the insertion validated at the blastocyst stage, the insertions accumulated in embryo-derived placenta are non-heritable LINE-1 insertions by definition. This is illustrated by our description of six LINE-1 insertions present at detectable levels only in the placenta, but not in blood DNA of newborns. Therefore, as lineage differentiation to epiblast/hypoblast or trophectoderm is known to occur at the morula stage (Niakan et al., 2012), we speculate that these *de novo* L1 insertions occurred in the polar trophectoderm during early embryogenesis, explaining their detection is such a large and complex tissue. However, we cannot exclude that some might have accumulated later during pregnancy.

It is worth noting that all *de novo* LINE-1 insertions uncovered in this study (n=16) are full-length or nearly full length (**Figure 7d**). Full-length elements can potentially undergo additional rounds of mobilization if they escape epigenetic silencing, further increasing the fluidity of pluripotent cells (Garcia-Perez et al., 2010). In addition, full-length insertions tend to be more mutagenic than 5’ truncated ones (reviewed in (Garcia-Perez et al., 2016; Goodier and Kazazian, 2008)). Previous studies in tumors and cultured tumor cells revealed that 5’-truncated insertions represent the most common outcome of LINE-1 retrotransposition (Carreira et al., 2013; Gilbert et al., 2005; Iskow et al., 2010; Lee et al., 2012; Tubio et al., 2014). In this study, the robust detection of 5’-truncated and full-length L1Hs elements annotated in the reference human genome, argue against a technical bias in the identification of *de novo* 5’-truncated insertions. Additionally, in this study we characterized and validated LINE-1 insertions with long polyA tails (even longer than 170-bp; average size polyA tails = 79-bp), even if sequencing homopolymeric sequences with Illumina can be challenging (Schirmer et al., 2016); thus, these data suggest that the presence of a long polyA tails does not affect our capability to detect L1-3’ ends and 5’-truncated insertions. Indeed, we confirmed robust detection and validation of 5’-truncated LINE-1 insertions in lung cancer samples using the same RC-seq protocol, despite harboring long polyA tails (average size polyA tails = 89-bp, n=10; **Figure S19**). A previous study analyzing LINE-1 retrotransposition in hiPSCs also revealed a high fraction of full-length insertions accumulated in these cells (Klawitter et al., 2016). The mechanism leading to 5’-truncation is not fully understood, but rather than reflecting an intrinsic limitation of the processivity of the LINE-1 Reverse Transcriptase (Gilbert et al., 2005; Monot et al., 2013; Piskareva and Schmatchenko, 2006), it seems linked to DNA-damage signaling and DNA repair pathways (Coufal et al., 2011; Suzuk i et al., 2009). Thus, it is possible that differences in the cellular milieu among cancer and pluripotent cells might affect 5’-truncation of new LINE-1 insertions. Less than 20% of all L1Hs elements in an average human genome are full-length (Myers et al., 2002). It is tempting to speculate that during embryonic development and/or in the germline, there might be negative selection mechanisms against the accumulation of full-length insertions (Boissinot et al., 2001; Boissinot et al., 2004), explaining this apparent discrepancy of results. Further genomic population studies will help clarifying this in the future.

In conclusion, in this study we have dissected LINE-1 expression and retrotransposition during early stages of human embryogenesis, revealing that L1-driven genetic heterogeneity starts to accumulate soon after fertilization, impacting the genome of cells that will give rise to the human body, but also occurs in extraembryonic lineages such as the embryo-derived placenta.

## METHODS SUMMARY

### Human embryonic stem cell culture and manipulation

hESCs were grown as previously described (Garcia-Perez et al., 2007; Macia et al., 2011). To establish clonal cell lines from hESCs, we used an engineered LINE-1 retrotransposition assay as described (Garcia-Perez et al., 2007).

### Human embryo manipulation

The whole procedure was approved by local regulatory authorities and the Spanish National Embryo steering committee. The embryos used in this study were of the highest quality possible. This is because Spanish legislation allows research on human embryos donated by couples who have undergone in vitro fertilization (IVF) (Cortes et al., 2007), and these were the embryos used in this study. Briefly, cryopreserved human embryos were anonymously donated with informed consent by couples that had already undergone an IVF cycle. All extractions/manipulations were carried out in a GMP certified facility (former Banco Andaluz Celulas Madre, Granada, Spain).

### LINE-1 expression analyses

LINE-1 mRNA expression was examined as described by Wissing et al. (Wissing et al., 2012), Coufal et al. (Coufal et al., 2009) and Macia et al. (Macia et al., 2011), with minor modifications. LINE-1 ORF1p expression was analyzed using a Zeiss LSM 710 confocal microscope.

### Whole Genome Amplification (WGA)

WGA was performed within 48 hours after single-cell extraction as described by Spit et al. (Spits et al., 2006). Negative controls of all the reagents used during the isolation of single cells were included (water, culture media, washing buffer, etc). The entire WGA procedure was performed in a class II laminar flow hood decontaminated by UV exposure for at least 30 min prior to use, in a GMP certified facility.

### RC-seq method

Multiplexed Illumina libraries were prepared from the different genomic DNAs and WGA-gDNA samples and pooled. LINE-1-insertion enrichment from libraries was performed using two biotinylated LNA-probes matching the 5’- and 3’-ends of the consensus sequence of a L1Hs-Ta element, as described (Sanchez-Luque et al., 2016; Upton et al., 2015). Captured libraries were paired-end sequenced (2×150bp reads) with an Illumina HiSeq2500/NextSeq500 platform, and computationally analyzed to discriminate *de novo* LINE-1 insertions from potential artifacts or previously reported/annotated insertions.

### ATLAS-seq method

ATLAS-seq was performed as previously described (Badge et al., 2003; Philippe et al., 2016) with minor modifications. Briefly, WGA-gDNAs were fragmented into 1kb fragments by sonication, end-repaired and linker-ligated. Separate suppression PCR reactions amplify L1Hs-Ta 5’- and 3’-junctions and add sample-specific barcodes, as well as Ion Torrent sequencing adapters. Libraries were multiplexed two-by-two and each pool was sequenced using a 318 Chip with a 400bp sequencing kit. The bioinformatic analysis scheme was adapted for WGA sequencing.

### Validation of LINE-1 insertions by PCR and capillary DNA sequencing

The number of LINE-1 insertions recovered by RC-seq and ATLAS-seq in human pre-implantation embryos, and analyzed by standard procedures, could not be directly compared due to inherent differences in sequencing technologies and L1 calling algorithms. ATLAS-seq uses clusters of single-end reads of variable length and sequencing/amplification artifacts are eliminated throughout the peak calling process. In contrast, RC-seq relies on paired-end reads of fixed length, and putative L1 insertions are primarily called for 1 read or few reads. As a consequence, the number of insertions detected by RC-seq in the absence of filtering was much higher than for ATLAS-seq (**Figure S11, S12 and S18** and Methods). To characterize LINE-1 insertions in hESCs and extraembryonic tissues, and as we used HMW-gDNAs, we only performed RC-seq without specific filtering analysis. All validations for putative L1 insertions were done with the same aliquot of gDNA or WGA-gDNA used in RC-seq and ATLAS-seq, using a class II laminar flow hood decontaminated by UV exposure for at least 30 min prior to use. Negative controls were included in all PCRs. We designed primers from the inferred insertion site avoiding repetitive areas of the genome. Hemi-nested 40+40 cycles PCRs were used for the 5’- and 3’-end filled-site PCRs. All amplification products were resolved on 1% agarose gels, excised, purified and fully sequenced using the Sanger method.

## ACKNOWLEDGEMENTS

We acknowledge current members of the J.L.G.-P. lab for helpful discussions. We also acknowledge Dr John V Moran (University of Michigan, US) for critical input during the project. The excellent technical support of Dr Purificación Catalina (karyotype analyses, BioBanco del SSPA, Granada), Ms Raquel Marrero (Microscopy Unit, Genyo) and the Genomic Unit of Genyo is also acknowledged. W e also thank Dr Oliver Weichenrieder (Max-Planck, Tubingen, Germany) for providing a polyclonal L1-ORF1p antibody. We are grateful to the IRCAN genomics core facility (GenoMed-UCA GenomiX) and C. Baudoin for Ion Torrent sequencing. The GenoMed facility is supported by FEDER, Conseil Départe mental 06, IBISA, Aviesan and INSERM. This project has been funded by a P lan de Estabilización de Grupos Emergentes de Investigación (ISCIII-CSJA-FEDER (EMER07/056)) and by a private donation to the lab from Ms Francisca Serrano (Trading y Bolsa para Torpes, Granada, Spain). Additionally, J.L.G.-P.’s lab is supported by CICE-FEDER-P12-CTS-2256, Plan Nacional de I+D+I 2008-2011 and 2013-2016 (FIS-FEDER-PI14/02152), PCIN-2014-115-ERA-NET NEURON II, the European Research Council (ERC-Consolidator ERC-STG-2012-233764), by an International Early Career Scientist grant from the Howard Hughes Medical Institute (IECS-55007420), and by The Wellcome Trust-University of Edinburgh Institutional Strategic Support Fund (ISFF2). S.R.H. is founded by Ramon y Cajal programme (RYC-2016-21395). R.V. is supported by the Spanish Ministry of Economy, Industry and Competitiveness (PFIS-FI16/00413) and this paper is part of her thesis project. D.C. was supported by the Spanish Ministry of Economy, Industry and Competitiveness (FPU-AP2010-0135). R.B. lab is funded by the University of Leicester. A.D.E. acknowledges the support of an ARC Discovery Early Career Researcher Award (DE150101117). G.J.F. acknowledges the support of CSL Centenary Fellowship. F.J.S.-L. is supported by the People Programme (Marie Curie Actions) of the European Union’s Seventh Framework Programme (FP7/2007-2013) under REA grant agreement n° PIOF-GA-2013-623324. Work in the laboratory of G.C. was supported by the French Government (National Research Agency, ANR) through the ‘Investments for the Future’ (LABEX SIGNALIFE, n°ANR-11-LABX-0028-01), by the Fondation pour la Recherche Médicale (FRM DEP20131128533), by the University Hospital Federation OncoAge, and by CNRS (GDR 3546).

## AUTHOR CONTRIBUTION

M.M.-L., R.B., G.J.F., G.C. and J.L.G.-P. conceived, designed, and interpreted the experiments. M.M.-L., C.P., R.R., R.V., T.W., S.R.R., J.L.C., E. G., D.C., S.R.H., F.J.S.-L., M.M., E.A., M.A.A., M.G.-C. and L.S. performed the experiments and data analyses. A.R., A.D.E. and G.C. assisted with the bioinformatic analyses. P.V., M.C.N-P, M.L., A.G.-M., B.G.-A. and C.A.-G. provided placenta and blood samples. S.A. provided expertise on human embryo donation and manipulation. M.M.-L., and J.L.G.-P. supervised the project. The manuscript was co-written by M.M.-L., G.C. and J.L.G.-P. with comments provided by all authors.

## COMPETING INTERESTS STATEMENT

The authors declare that they have no competing financial interests.

**Supplementary Information** is linked to the online version of the paper.

